# Medial septum parvalbumin-expressing inhibitory neurons are impaired in a mouse model of Dravet Syndrome

**DOI:** 10.1101/2024.10.29.620933

**Authors:** Limei Zhu, Yiannos Demetriou, Joseph Barden, Juan Disla, Joanna Mattis

**Affiliations:** Department of Neurology, University of Michigan, Ann Arbor, MI USA; M.D./Ph.D. Program, Wayne State University, Detroit, MI USA; Michigan Neuroscience Institute, University of Michigan, Ann Arbor, MI USA

**Keywords:** Dravet Syndrome, epilepsy, medial septum, parvalbumin

## Abstract

Dravet syndrome (DS) is a severe neurodevelopmental disorder caused by pathogenic variants in the *SCN1A* gene, which encodes the voltage-gated sodium channel Na_v_1.1 α subunit. Experiments in animal models of DS – including the haploinsufficient *Scn1a*^+/-^ mouse – have identified impaired excitability of interneurons in the hippocampus and neocortex; this is thought to underlie the treatment-resistant epilepsy that is a prominent feature of the DS phenotype. However, additional brain structures, such as the medial septum (MS), also express *SCN1A*. The medial septum is known to play an important role in cognitive function and thus may contribute to the intellectual impairment that also characterizes DS. In this study, we employed whole cell patch clamp recordings in acute brain slices to characterize the electrophysiological properties of MS neurons in *Scn1a*^+/-^ mice versus age-matched wild-type littermate controls. We found no discernible genotype-related differences in MS cholinergic (ChAT) neurons, but significant dysfunction within MS parvalbumin-expressing (PV) inhibitory neurons in *Scn1a*^+/-^ mice. We further identified heterogeneity of firing patterns among MS PV neurons, and additional genotype differences in the proportion of subtype representation. These results confirm that the MS is an additional locus of pathology in DS, that may contribute to co- morbidities such as cognitive impairment.

## Introduction

Dravet syndrome (DS) is an infantile-onset developmental and epileptic encephalopathy characterized by treatment-resistant epilepsy, temperature-sensitive seizures, developmental delay, intellectual disability, features of autism spectrum disorder, and an increased risk of sudden unexpected death in epilepsy (SUDEP) (Villas et al., 2017). Most patients with DS harbor pathogenic loss-of-function variants in the *SCN1A* gene, leading to effective haploinsufficiency of the voltage-gated sodium channel Na_v_1.1 α subunit (Claes et al., 2001).

The hemizygous *Scn1a*^+/-^ mouse (Mistry et al., 2014) is a well-established preclinical model that accurately replicates key phenotypic features of the human condition. Studies using this and other DS models have provided crucial insights into cellular pathology of the disorder. Notably, all major classes of GABAergic neurons have been found to exhibit impaired action potential generation in mouse models of DS (e.g., (Favero et al., 2018; Goff & Goldberg, 2019; Mattis et al., 2022; Rubinstein et al., 2015; Tai et al., 2014; Tsai et al., 2015)).

Most pre-clinical data characterizing DS pathology has focused on seizure-prone regions, such as the hippocampus and neocortex. However, in addition to seizures, patients with DS also have prominent cognitive, motor, and sleep dysfunction (Darra et al., 2019; Licheni et al., 2018; Selvarajah et al., 2022; Wolff et al., 2006). Even as seizure burden tends to decrease over time, these other symptoms persist and significantly impact quality of life (Villas et al., 2017). As the expression of *SCN1A* is brain-wide, there is therefore a need to better understand the pathology of DS in other brain regions that may contribute to these diverse symptoms.

Hippocampal theta oscillations provide temporal organization for neuronal assemblies, with relevance for learning and memory (Buzsáki, 2002; Buzsáki & Moser, 2013; Colgin, 2013). Theta oscillations are controlled by the medial septum (MS), a cholinergic nucleus in the basal forebrain that sends dense projections to the hippocampus (Freund & Antal, 1988; Huh et al., 2010; Lee et al., 1994; Vandecasteele et al., 2014). A significant majority of cholinergic neurons and nearly all GABAergic neurons in the MS are reported to express Na_v_1.1 (Bender et al., 2016). Furthermore, abnormalities of the hippocampal theta rhythm have been identified in animal models of DS (Bender et al., 2013, 2016; Jansen et al., 2021), which may contribute to cognitive impairment.

We hypothesized that MS neurons have impaired excitability in DS. To test this, we employed acute slice whole cell patch-clamp recordings to compare the electrophysiological characteristics of cholinergic (ChAT) and inhibitory, parvalbumin-expressing (PV) neurons in *Scn1a*^+/-^ mice versus age-matched wild-type (WT) littermate controls. Our findings revealed no discernible genotype-related differences in MS ChAT neurons; however, we observed a pronounced hypoexcitability across all subtypes of PV neurons in *Scn1a*^+/-^ mice. These findings support the concept that the MS is a locus of pathology in DS and may contribute to intellectual disability and other comorbidities.

## Results

As Na_v_1.1 is reportedly expressed in approximately 90% of cholinergic neurons in the MS (Bender et al., 2016), we began by evaluating the excitability of choline acetyltransferase-expressing (ChAT) MS neurons from *Scn1a*^+/-^ mice compared to age-matched, WT littermate controls. To enable targeted patching of ChAT neurons in slice, we crossed ChAT-Cre and tdTomato (tdT) reporter mice (ChAT-Cre.TdT), which we bred to double-homozygosity on a C57BL/6 background. We then further crossed these mice with *Scn1a*^+/-^ mice, which were maintained on a 129S6.SvEvTac background. All experimental F1 mice (*Scn1a*^*+/-*^.ChAT-Cre.TdT and *Scn1a*^*+/+*^. ChAT-Cre.TdT) were therefore on the 50:50 129S6:C57BL/6J genetic background that is standard in the field (Mistry et al., 2014) (**Figure 1A**). We next confirmed that tdT is selectively expressed in MS ChAT neurons by immunostaining for ChAT and quantifying the co- localization. We found a substantial overlap of tdT expression and ChAT immunopositivity: 99% of tdT-positive neurons stained for ChAT, and 73% of ChAT neurons were well labeled by tdT (**Figure 1B-C**).

**Figure 1.**
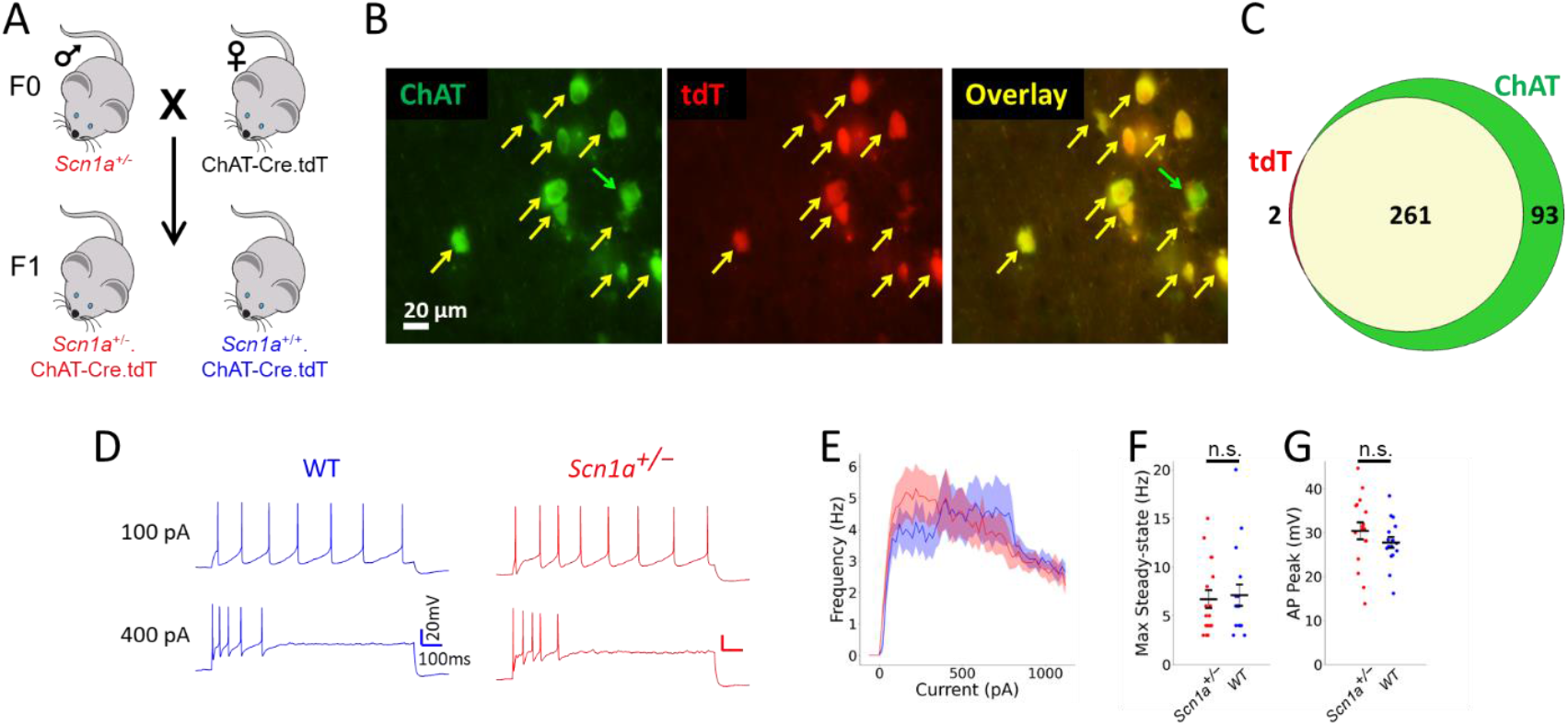
MS ChAT neurons have similar firing properties in *Scn1a*^*+/-*^ and WT mice. **(A)** Strategy to generate mice for targeted patching of cholinergic neurons. Female ChAT-Cre.tdT double-homozygous mice are bred with male *Scn1a*^*+/-*^ mice. The F1 progeny include *Scn1a*^*+/-*^.ChAT-Cre.tdT and *Scn1a*^*+/+*^.ChAT-Cre.tdT mice that have tdT expression in ChAT cells. **(B)** Representative images of neurons with choline acetyltransferase (ChAT) immunostaining (green; left) and tdT-expressing neurons (red; middle). The merged image (right) illustrates substantial overlap between tdT and ChAT expression (yellow arrows), with rare neurons that immunostain for ChAT but do not express tdT (green arrow). Scale bars 20 μm. **(C)** Venn diagram showing the quantification of medial septum neurons that immunostain for ChAT and/or express tdT. Of the 263 neurons that were tdT-positive, 261 (99%) immunostained for ChAT. *n* = 356 neurons across 4 mice, P45-92. **(D)** Example current clamp recordings of medial septum ChAT cells from WT (blue) and *Scn1a*^*+/-*^ (red) mice, showing action potentials evoked by 100 and 400 pA current injections. **(E)** No difference was observed in the current/frequency (I-f) plot. Line and shaded areas represent mean and SEM; 2-way ANOVA. Maximal steady state firing frequency **(F)** and AP peak **(G)** were similar across genotypes; unpaired two-way t-test. n = 17 cells / 5 mice for WT and 17 cells / 6 mice for *Scn1a*^*+/-*^.

We prepared acute brain slices from these triple-transgenic mice at postnatal day (P)21- 30 and performed targeted whole-cell patch-clamp recordings from tdT+ (i.e., ChAT) neurons in the MS. We found that MS ChAT neurons were slow-firing, consistent with previous reports (Griffith, 1988; Griffith & Matthews, 1986; Markram & Segal, 1990; Mattis et al., 2014; Sotty et al., 2003; Yi et al., 2021) (**Figure 1D**). We identified no significant differences between the genotypes, as shown by similar input-output curves (**Figure 1D-E**) and quantification across multiple measurements of electrophysiological properties (**Figure 1F-G; Table 1**).

**Table 1.** *Scn1a*^*+/-*^ MS ChAT neurons have normal properties. Properties of MS ChAT neurons from *Scn1a*^*+/-*^ and WT mice. Significance is determined by unpaired two-way t-test, corrected for multiple comparisons within each group.

| Measurement | WT | *Scn1a*<sup>+/-</sup> | p-Value (corrected) |
| --- | --- | --- | --- |
| *n* cells (mice) | 17(5) | 17 (6) |  |
| **Intrinsic Properties** |  |  |  |
| V<sub>m</sub> (mV) | -58.6 ± 1.4 | -56.7 ± 0.9 | 0.54 |
| R<sub>m</sub> (MΩ) | 531 ± 38 | 679 ± 59 | 0.21 |
| Time Constant (ms) | 27.8 ± 2.3 | 25.7 ± 1.0 | 0.54 |
| Rheobase (pA) | 49 ± 6 | 39 ± 8 | 0.54 |
| **Repetitive AP Properties** |  |  |  |
| Max Steady-state (Hz) | 7.1 ± 1.1 | 6.7 ± 0.9 | 0.78 |
| Max Instantaneous (Hz) | 138.1 ± 8.6 | 122.2 ± 7.9 | 0.73 |
| **Individual AP Properties** |  |  |  |
| AP Threshold (mV) | -33.0 ± 0.6 | -31.9 ± 0.8 | 0.45 |
| AP Peak (mV) | 27.7 ± 1.2 | 30.4 ± 1.9 | 0.45 |
| AP Amplitude (mV) | 60.8 ± 1.3 | 62.3 ± 1.9 | 0.52 |
| AP Rise Time (ms) | 0.7 ± 0.0 | 0.8 ± 0.0 | 0.47 |
| APD50 (ms) | 1.5 ± 0.1 | 1.6 ± 0.1 | 0.34 |
| APD90 (ms) | 1.9 ± 0.1 | 2.2 ± 0.1 | 0.45 |

We hypothesized that deficits may be evident in non-cholinergic MS cell types, so we next compared the properties of tdT-negative (i.e., ChAT-negative) MS neurons in *Scn1a*^+/-^ versus WT control mice. Indeed, we found that MS ChAT-negative neurons in *Scn1a*^+/-^ mice were hypoexcitable, with reduced firing frequency (**Figure S1**). However, MS ChAT-negative neurons are known to be a heterogeneous mix of glutamatergic neurons and at least two types of GABAergic neurons, parvalbumin (PV)-expressing and somatostatin (SST)-expressing cells, which have distinct firing properties (Sotty et al., 2003; Yi et al., 2021). We therefore opted to selectively record from MS PV neurons, since nearly all GABAergic MS neurons reportedly express Na_v_1.1 (Bender et al., 2016), and since *Scn1a*^+/-^ PV neurons show pronounced deficits in hippocampus and neocortex (De Stasi et al., 2016; Favero et al., 2018; Mattis et al., 2022; Tai et al., 2014).

To selectively assay MS PV neurons, we employed the above-described breeding strategy to generate triple-transgenic mice in which tdT expression was under Cre-dependent control and expressed in PV neurons (**Figure 2A**). Quantification of tdT expression and PV immunostaining revealed 86% of tdT-expressing neurons were PV-positive, and 77% of PV neurons were successfully labeled by tdT (**Figure 2B-C**). We prepared acute brain slices at P21-30 and performed whole-cell current-clamp recordings from 101 fluorescently labeled MS PV neurons across *Scn1a*^+/-^ mice and age-matched WT littermate controls. We found that MS PV neurons from *Scn1a*^+/-^ mice exhibited multiple abnormalities (**Table 2**) including a marked decrease in repetitive action potential (AP) firing (**Figure 2D-F**) and significantly reduced AP peak (**Figure 2G**).

**Table 2.** *Scn1a*^*+/-*^ MS PV neurons have impaired properties. Properties of MS PV cells from *Scn1a*^*+/-*^ and WT mice. Significance is determined by unpaired two-way t-test, corrected for multiple comparisons within each group.

| Measurement | WT | *Scn1a*<sup>+/-</sup> | p-Value (corrected) |
| --- | --- | --- | --- |
| *n* cells (mice) | 49 (13) | 52 (11) |  |
| **Intrinsic Properties** |  |  |  |
| V<sub>m</sub> (mV) | -63.4 ± 0.6 | -64.3 ± 0.6 | 0.54 |
| R<sub>m</sub> (MΩ) | 192 ± 21 | 216 ± 22 | 0.54 |
| Time Constant (ms) | 9.1 ± 0.7 | 10.6 ± 0.8 | 0.54 |
| Rheobase (pA) | 83 ± 6 | 84 ± 7 | 0.82 |
| **Repetitive AP Properties** |  |  |  |
| Max Steady-state (Hz) | 145 ± 13 | 87 ± 9 | **0.002 (\*\*)** |
| Max Instantaneous (Hz) | 414 ± 9 | 384 ± 9 | **0.03 (\*)** |
| SFAn | 0.51 ± 0.02 | 0.45 ± 0.03 | 0.15 |
| **Individual AP Properties** |  |  |  |
| AP Threshold (mV) | -38.2 ± 0.7 | -38.2 ± 0.5 | 0.99 |
| AP Peak (mV) | 26.8 ± 0.9 | 22.4 ± 1.0 | **0.01 (\*)** |
| AP Amplitude (mV) | 65.1 ± 1.3 | 60.6 ± 1.0 | **0.02 (\*)** |
| AP Rise Time (ms) | 0.39 ± 0.01 | 0.42 ± 0.01 | 0.09 |
| APD50 (ms) | 0.62 ± 0.02 | 0.68 ± 0.01 | **0.03 (\*)** |
| APD90 (ms) | 0.77 ± 0.02 | 0.85 ± 0.02 | **0.03 (\*)** |

**Figure 2.**
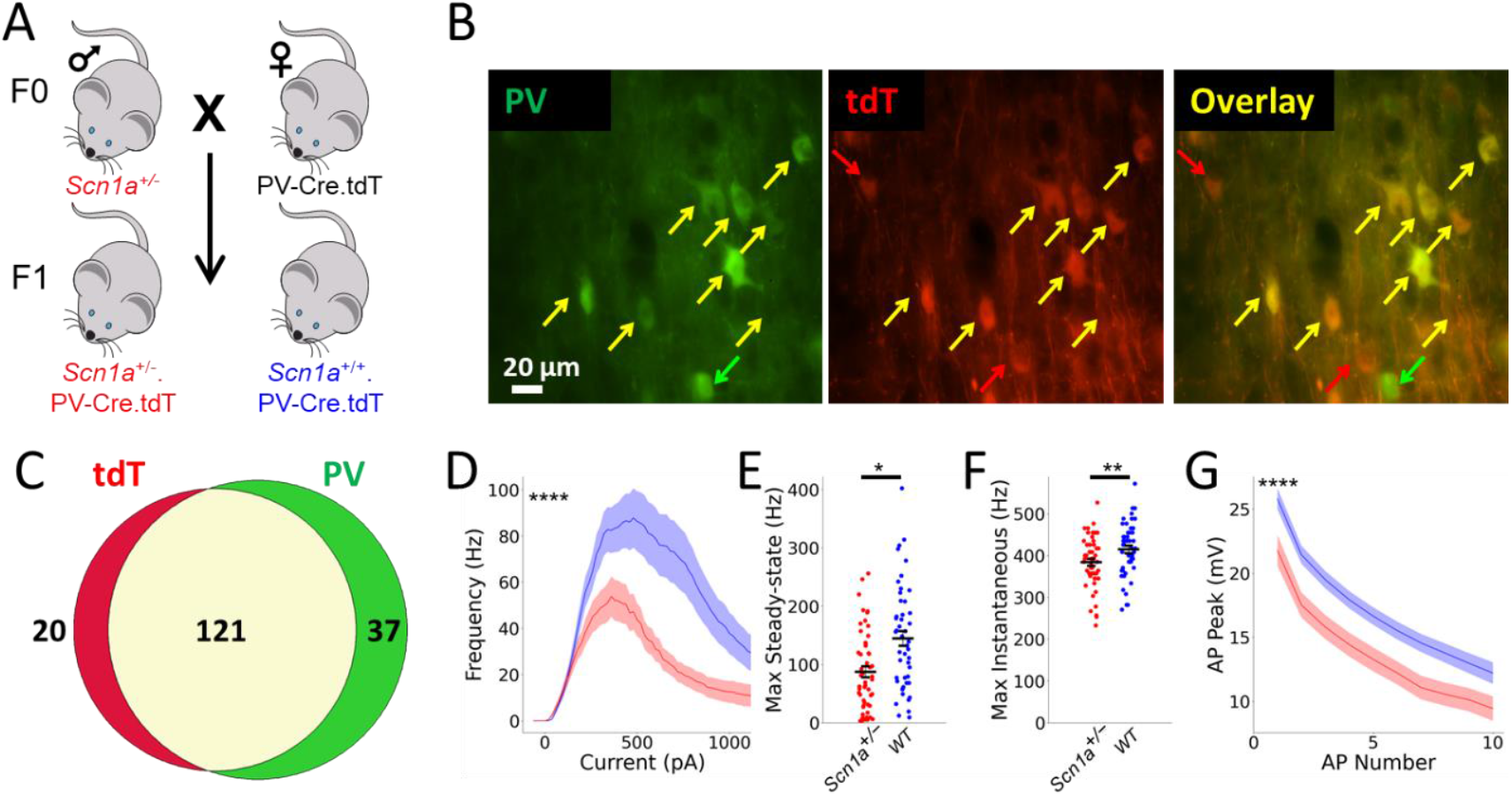
MS PV neurons have impaired firing in *Scn1a*^*+/-*^ mice. **(A)** Strategy to generate mice for targeted patching of parvalbumin-expressing neurons. Female PV-Cre.tdT double- homozygous mice are bred with male *Scn1a*^*+/-*^ mice. The F1 progeny include *Scn1a*^*+/-*^.PV- Cre.tdT and *Scn1a*^*+/+*^.PV-Cre.tdT mice that have tdT expression in PV cells. **(B)** Representative images of neurons with PV immunostaining (green; left) and tdT-expressing neurons (red; middle). The merged image (right) illustrates substantial overlap between tdT and PV expression (yellow arrows). Rare neurons express tdT but do not immunostain for PV (red arrow) or immunostain for PV but do not express tdT (green arrow). **(C)** Venn diagram showing the quantification of medial septum neurons that immunostain for PV and/or express tdT. **(D)** Current/frequency (I-f) plot of PV cells patched within medial septum demonstrates hypoexcitability of *Scn1a*^*+/-*^ cells. Both maximal steady-state **(E)** and instantaneous **(F)** firing frequencies were significantly lower for *Scn1a*^*+/-*^ cells. **(G)** AP number versus AP peak also demonstrated a significant difference between genotypes, with a more rapid decline *Scn1a*^*+/-*^ cells. n = 49 cells / 13 mice for WT and n = 52 cells / 11 mice for *Scn1a*^*+/-*^. For **D** and **G**, stars in the upper left indicate overall genotype significance (2-way ANOVA), and line and shaded areas represent mean and SEM. For **E** and **F**, stars indicate significance by unpaired t-test. (****) indicates p < 0.0001, (**) indicates p < 0.01, and (*) indicates p < 0.05.

We hypothesized that the hypoexcitability of MS PV neurons in *Scn1a*^*+/-*^ mice was due to insufficient Na_v_1.1 expression. To test this, we examined the impact on these cells of Hm1a, a spider venom toxin that selectively potentiates Na_v_1.1 by preventing its inactivation (Osteen et al., 2016, 2017). Indeed, we found that bath-application of Hm1a significantly increased firing of *Scn1a*^*+/-*^ MS PV neurons (**Figure 3**).

**Figure 3.**
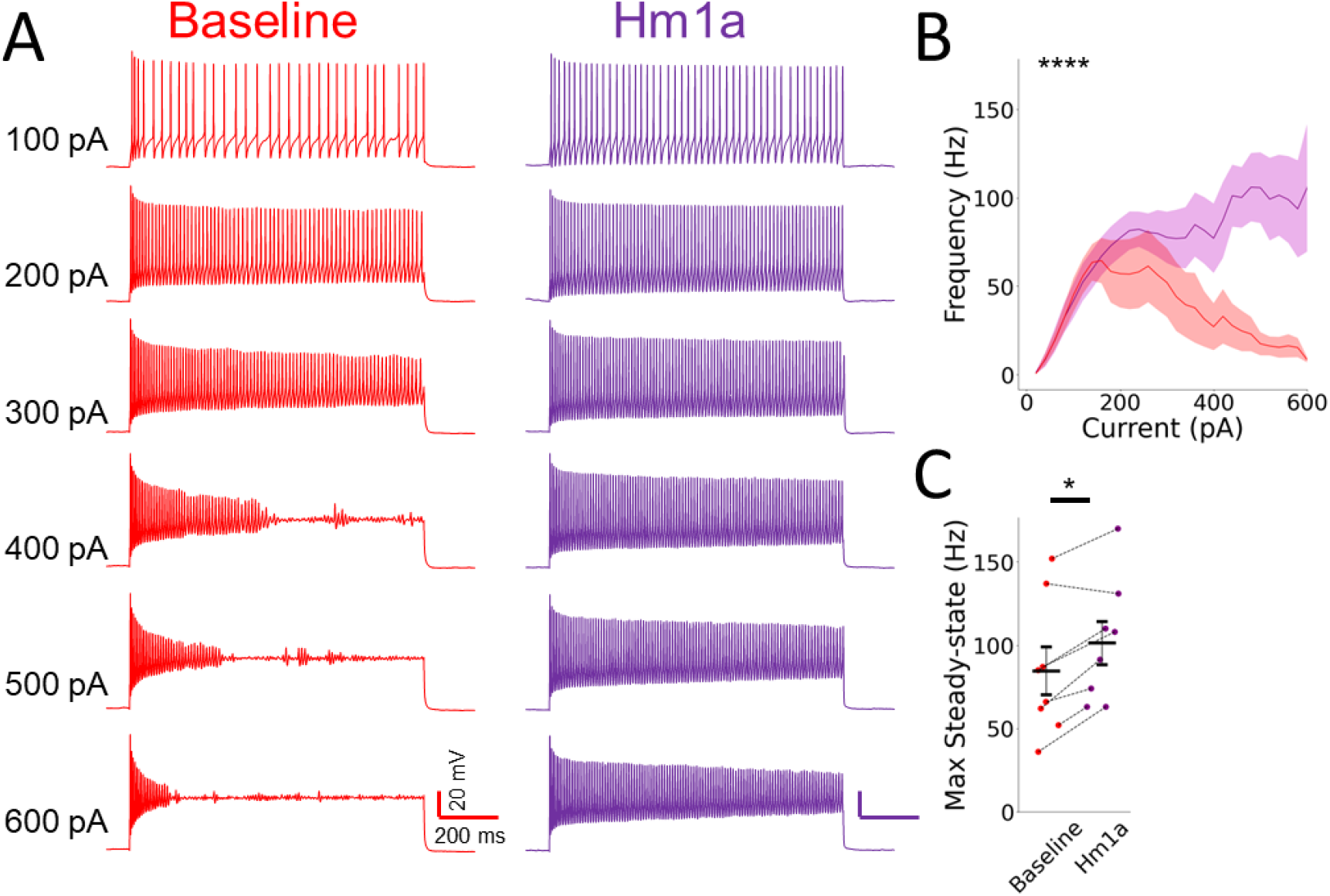
Bath-application of Hm1a increases firing of MS PV neurons in *Scn1a*^*+/-*^ mice. **(A)** Representative current clamp recordings from an *Scn1a*^*+/-*^ MS PV neuron before and after bath-application of Hm1a. **(B)** Current/frequency (I-f) plot shows significantly increased firing with larger current injections after Hm1a application (purple) relative to baseline (red). Line and shaded areas represent mean and SEM; 2-way ANOVA. **(C)** Maximal steady-state firing frequencies significantly increased after Hm1a application, as determined by paired t-test. (****) indicates p < 0.0001 and (*) indicates p < 0.05.

We additionally examined whether the difference in MS PV neuronal physiology could be due to abnormalities in MS perineuronal nets (PNNs), which form a lattice of extracellular matrix components around neurons, particularly PV neurons, including in the MS (Morris & Henderson, 2000), and may be disrupted in epilepsy (Chaunsali et al., 2021). However, we detected no genotype difference in the intensity of fluorescence signal of labeled PNNs within the MS (**Figure S2**).

As illustrated in **Figure 4A-C**, we observed that MS PV neurons in both WT and *Scn1a*^+/-^ mice exhibited several distinct firing patterns in response to depolarizing step currents. We used two parameters to separate these subtypes – burst length (i.e., the length of the maximal burst of continuous firing), and the coefficient of variation of the inter-spike interval (ISI CoV) – both of which were calculated from the current step trace in which a cell fired the maximal number of APs (**Figure 4D**). We defined “fast-firing” neurons as those with sustained firing throughout the pulse train (burst length > 800 ms for a 1000 ms current step); these all had a low ISI CoV. We defined “burst-firing” neurons as those with a shorter burst length (< 800 ms) and low ISI CoV (<1); these invariably fired their burst at the onset of the pulse. Finally, we defined “cluster-firing” neurons – which fired in clusters of spikes throughout the pulse – as having a high ISI CoV (>1).

**Figure 4.**
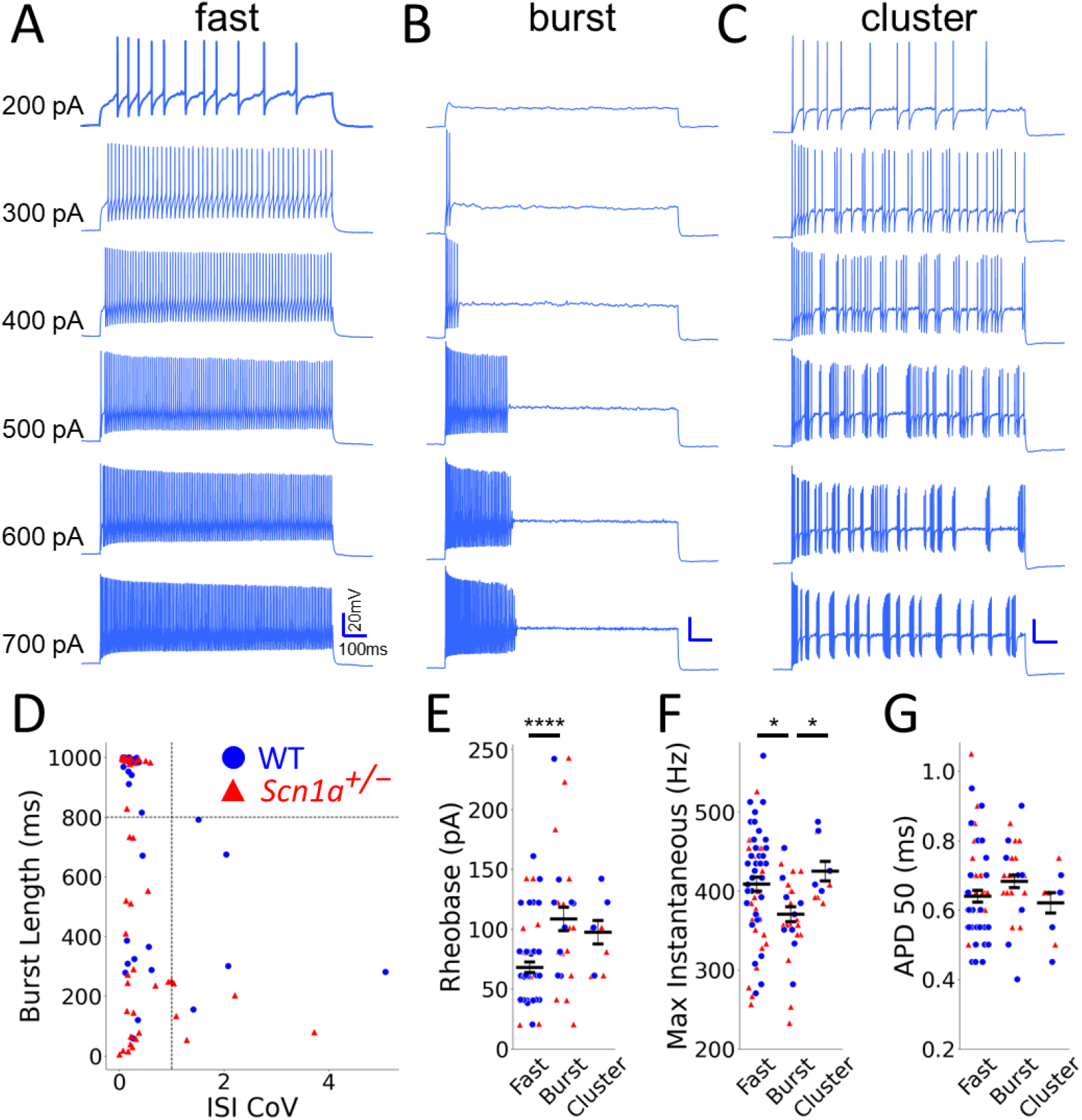
Diversity of firing properties within PV MS neurons. Representative current clamp recordings of WT MS PV cells with **(A)** fast-firing, **(B)** burst-firing, and **(C)** cluster-firing patterns. **(D)** Cells were categorized based on the length of the initial burst (burst length) and the coefficient of variation of the inter-spike interval (ISI CoV). Dashed lines indicate cut-off values used to define groups. Cells with a low ISI CoV and long burst length were defined as fast-firing. Cells with shorter burst length were defined as burst-firing (low ISI CoV) or cluster-firing (high ISI CoV). **(E)** Fast-firing cells have the lowest rheobase (68 ± 4 pA, versus 109 ± 10 pA for burst-firing and 97 ± 10 pA for cluster-firing). All PV subtypes displayed a high maximal instantaneous firing frequency (ranging from 370 ± 9 Hz for burst-firing to 425 ± 13 Hz for cluster-firing) **(F)** and brief action potential duration (ranging from 0.62 ± 0.03 ms for cluster- firing to 0.68 ± 0.02 ms for burst-firing) **(G)**. n = 60 cells / 24 mice (fast-firing), 31 cells / 14 mice (burst-firing), and 10 cells / 9 mice (cluster-firing). Stars indicate significance by unpaired t-test. (*) indicates p < 0.05 and (****) indicates p < 0.0001.

We compared these subtypes of MS PV neurons across a wide range of additional parameters (**Table 3**). As a direct reflection of our clustering strategy, there were highly significant differences across subtypes in burst length (as well as maximal steady state firing frequency) and ISI CoV. We additionally found significant subtype variability in rheobase, with the fast-firing group having the lowest (**Figure 4E**), suggesting these were more readily excitable. However, all subtypes displayed characteristic properties of PV cells, with high maximal instantaneous firing rate (**Figure 4F**) and narrow spikes (**Figure 4G**).

**Table 3.** Properties of all MS PV cell subtypes in WT and *Scn1a*^*+/-*^ mice. Properties of fast-firing, burst-firing, and cluster-firing MS PV cells from *Scn1a*^*+/-*^ and WT mice. Significance is determined by two-way ANOVA.

|  | Fast-firing |  | Burst-firing |  | Cluster-firing |  | 2-way ANOVA |  |
| --- | --- | --- | --- | --- | --- | --- | --- | --- |
| Measurement | WT | *Scn1a*<sup>+/-</sup> | WT | *Scn1a*<sup>+/-</sup> | WT | *Scn1a*<sup>+/-</sup> | p-Value (cell type) | p-Value (genotype) |
| *n* cells (mice) | 35 (13) | 25 (11) | 9 (5) | 22 (9) | 5 (4) | 5 (5) |  |  |
| Intrinsic Properties |  |  |  |  |  |  |  |  |
| V<sub>m</sub> (mV) | -63.4 ± 0.8 | -64.0 ± 1.0 | -64.0 ± 0.8 | -64.4 ± 0.6 | -61.7 ± 1.7 | -65.7 ± 0.8 | 0.88 | 0.14 |
| R<sub>m</sub> (MΩ) | 179 ± 13 | 256 ± 40 | 266 ± 100 | 189 ± 25 | 146 ± 20 | 143 ± 10 | 0.33 | 0.96 |
| Time Constant (ms) | 9.5 ± 0.9 | 12.8 ± 0.9 | 9.8 ± 1.9 | 8.9 ± 1.4 | 6.1 ± 0.3 | 7.5 ± 1.0 | **0.02 (\*)** | 0.34 |
| Rheobase (pA) | 70 ± 6 | 65 ± 6 | 117 ± 18 | 105 ± 12 | 105 ± 13 | 89 ± 15 | **<0.0001 (\*\*\*\*)** | 0.31 |
| Repetitive AP Properties |  |  |  |  |  |  |  |  |
| Max Steady-state (Hz) | 177 ± 14 | 142 ± 11 | 46 ± 11 | 31 ± 5 | 96 ± 22 | 57 ± 16 | **<0.0001 (\*\*\*\*)** | 0.06 |
| Max Instantaneous (Hz) | 422 ± 11 | 390 ± 14 | 371 ± 17 | 370 ± 12 | 440 ± 18 | 411 ± 17 | **0.02 (\*)** | 0.21 |
| SFAn | 0.53 ± 0.02 | 0.51 ± 0.03 | 0.44 ± 0.05 | 0.42 ± 0.05 | 0.50 ± 0.11 | 0.25 ± 0.14 | **0.02 (\*)** | **0.04 (\*)** |
| Burst length (ms) | 984 ± 6 | 987 ± 7 | 311 ± 58 | 255 ± 49 | 441 ± 123 | 143 ± 36 | **<0.0001 (\*\*\*\*)** | **0.001 (\*\*)** |
| ISI CoV | 0.15 ± 0.01 | 0.19 ± 0.03 | 0.32 ± 0.06 | 0.30 ± 0.06 | 2.42 ± 0.67 | 1.87 ± 0.51 | **<0.0001 (\*\*\*\*)** | **0.002 (\*\*)** |
| Individual AP Properties |  |  |  |  |  |  |  |  |
| AP Threshold (mV) | -39.0 ± 0.8 | -38.4 ± 0.7 | -35.2 ± 1.5 | -37.6 ± 1.0 | -38.3 ± 1.6 | -40.3 ± 0.5 | 0.06 | 0.27 |
| AP Peak (mV) | 27.4 ± 1.1 | 23.6 ± 1.5 | 25.0 ± 2.3 | 22.2 ± 1.4 | 26.2 ± 2.7 | 17.7 ± 2.4 | 0.22 | **0.006 (\*\*)** |
| AP Amplitude (mV) | 66.4 ± 1.6 | 62.0 ± 1.6 | 60.2 ± 2.4 | 59.8 ± 1.3 | 64.5 ± 1.8 | 57.9 ± 2.3 | 0.07 | 0.07 |
| AP Rise Time (ms) | 0.38 ± 0.01 | 0.42 ± 0.02 | 0.43 ± 0.04 | 0.42 ± 0.01 | 0.39 ± 0.03 | 0.38 ± 0.03 | 0.19 | 0.71 |
| APD50 (ms) | 0.61 ± 0.02 | 0.68 ± 0.03 | 0.67 ± 0.05 | 0.69 ± 0.02 | 0.60 ± 0.04 | 0.64 ± 0.04 | 0.32 | 0.20 |
| APD90 (ms) | 0.76 ± 0.03 | 0.84 ± 0.03 | 0.81 ± 0.06 | 0.86 ± 0.02 | 0.75 ± 0.06 | 0.80 ± 0.06 | 0.45 | 0.16 |

We next evaluated whether the impairment observed in the combined pool of *Scn1a*^+/-^ MS PV cells (**Figure 2**) was seen across each of these different subtypes. Indeed, we noted decreased firing of *Scn1a*^+/-^ MS fast-firing, burst-firing, and cluster-firing PV cells (**Figure 5A-C**). Quantitatively, all *Scn1a*^+/-^ MS PV cells subtypes had significantly impaired repetitive firing that emerged in response to larger depolarizing current injections (**Figure 5D, F, H**). We again found significantly reduced AP peak (**Table 3**) and pronounced spike run-down (**Figure 5E, G, I**) in *Scn1a*^+/-^ cells relative to WT controls. There were no significant genotype differences in intrinsic neuron properties, including resting potential, input resistance, membrane time constant, or rheobase (**Table 3**).

**Figure 5.**
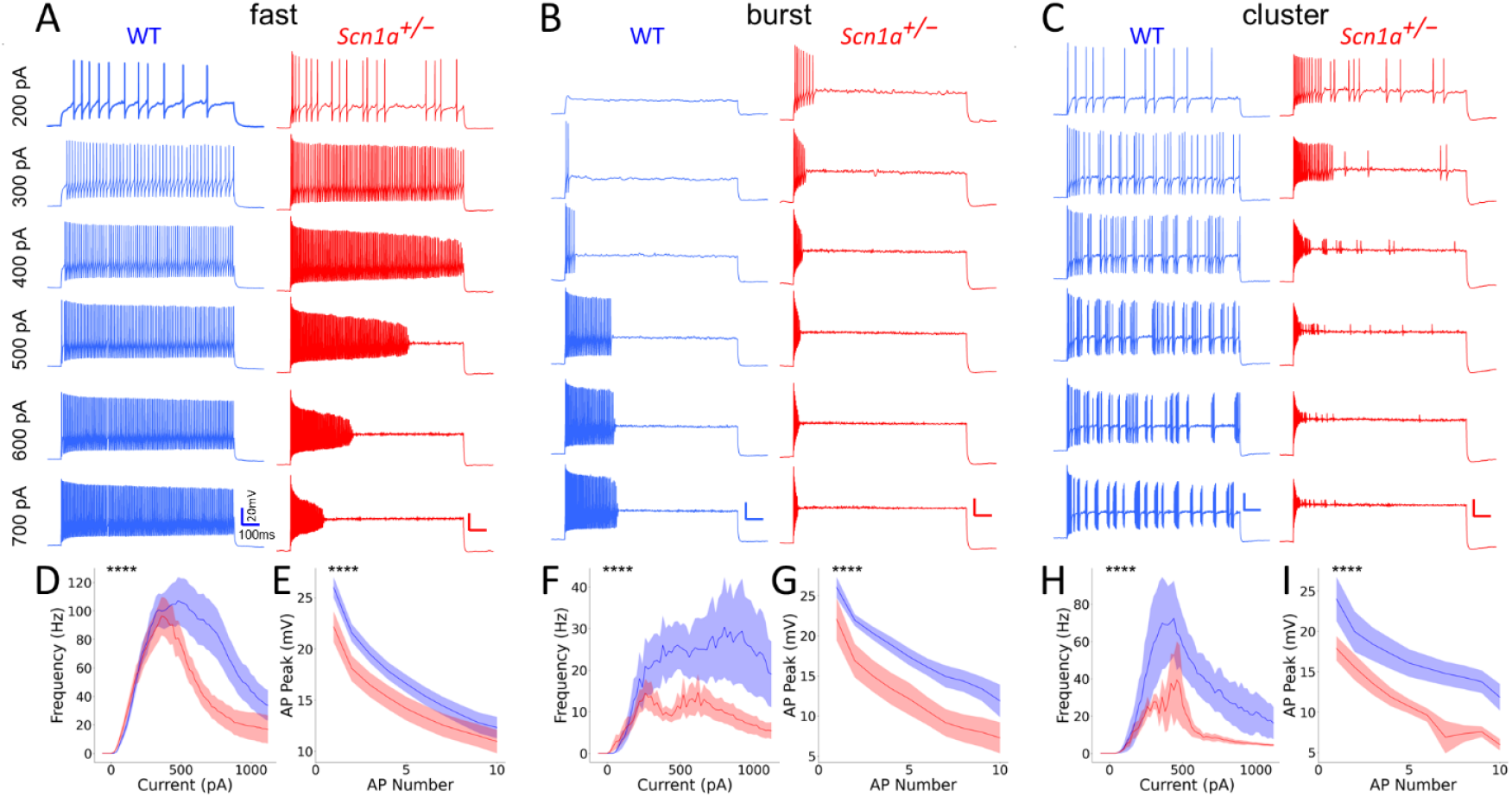
Impaired spike generation is seen across all subtypes of MS PV cells in *Scn1a*^*+/-*^ mice. Representative current clamp recordings of MS PV cells classified as fast-firing **(A)**, burst-firing **(B)**, and cluster-firing **(C)** from WT (blue) and *Scn1a*^*+/-*^ (red) mice. These traces illustrate progressively impaired spike generation in *Scn1a*^*+/-*^ cells in response to depolarizing current injections. Fast-firing *Scn1a*^*+/-*^ cells displayed significantly impaired action potential firing **(D)** and decreased AP peak **(E)** relative to fast-firing WT cells (n = 35 cells / 13 mice for WT and 25 cells / 11 mice for *Scn1a*^*+/-*^.) Burst-firing *Scn1a*^*+/-*^ cells displayed significantly impaired action potential firing **(F)** and decreased AP peak **(G)** relative to burst-firing WT cells (n = 9 cells / 5 mice for WT and 22 cells / 9 mice for *Scn1a*^*+/-*^.). Cluster-firing *Scn1a*^*+/-*^ cells displayed significantly impaired action potential firing **(H)** and decreased AP peak **(I)** relative to cluster-firing WT cells (n = 5 cells / 4 mice for WT and 5 cells / 5 mice for *Scn1a*^*+/-*^). Line and shaded areas represent mean and SEM. For **D-I**, stars in the upper left reflect overall genotype comparisons, which were determined via 2-way ANOVA. (****) indicates p < 0.0001.

Finally, we asked whether there was a genotype difference not only in the properties of neurons within each MS PV subtype, but also in the relative proportion of the subtypes (**Figure 6**). Indeed, the burst-firing subtype constituted only ∼18% of MS PV cells in WT mice but was more than double that (∼42%) in *Scn1a*^+/-^ mice. Conversely fast-firing PV cells were the large majority (∼72%) encountered in WT mice but less than half (∼48%) in *Scn1a*^+/-^ mice. Cluster- firing cells constituted ∼10% of the population in both genotypes.

**Figure 6.**
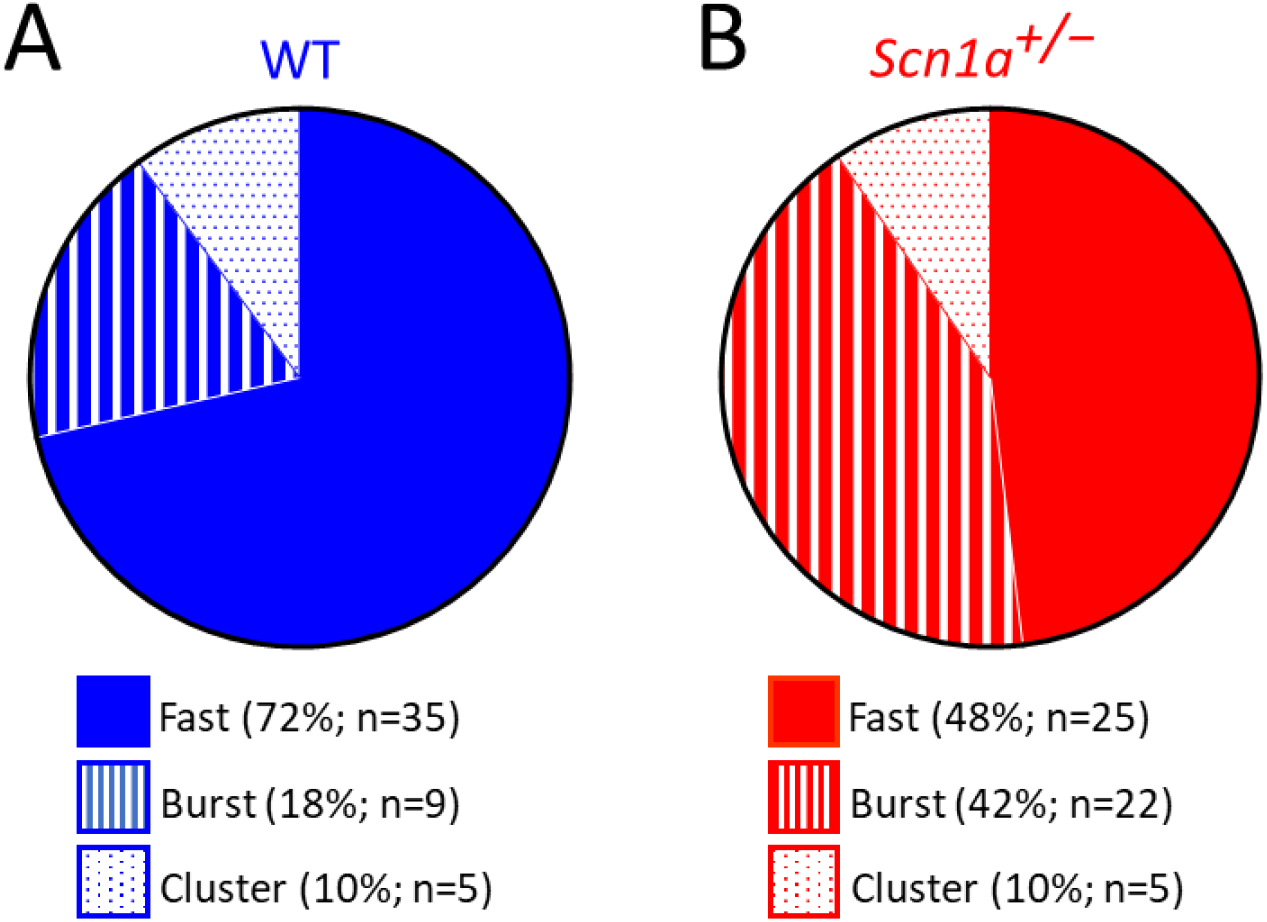
Genotype differences in proportion of different MS PV neurons. *Scn1a*^*+/-*^ mice have a greater proportion of medial septum burst-firing PV cells, relative to WT littermate control mice. Proportion of WT **(A)** and *Scn1a*^+/-^ **(B)** MS PV cells that exhibited continuous (solid), burst (striped), or cluster (dotted) firing patterns. The *Scn1a*^+/-^ PV cell population was highly enriched for burst-firing cells relative to WT (42% vs 18% burst cells; p = 0.002; Chi-square test).

## Discussion

Prior work has identified impairment of PV neurons in mouse models of DS. However, these studies have mainly focused on neurons of the neocortex and hippocampus – i.e., regions that are known to be highly ictogenic. Subcortical brain regions may also contribute to the DS phenotype, but the impact of Na_v_1.1 haploinsufficiency on these regions remains largely unknown. In this study, we performed targeted patching of ChAT and PV neurons in the MS of *Scn1a*^*+/-*^ and age-matched, WT littermate control mice. We found that although MS ChAT neurons are spared, MS PV neurons are significantly hypoexcitable in *Scn1a*^+/-^ mice, across all identified subtypes as defined by firing pattern. These findings may implicate a medial septal contribution to the cognitive impairment that is a prominent feature of DS.

### MS ChAT neurons are functionally preserved in *Scn1a*^*+/-*^ mice

Although approximately 90% of MS ChAT neurons are reported to express Na_v_1.1 (Bender et al., 2016), we identified no differences in these neurons – including no changes in cell intrinsic properties, measurements of repetitive firing, or features of individual action potentials – in *Scn1a*^+/-^ mice relative to WT littermate controls (**Figure 1**).

There are multiple potential explanations for this negative result. First, Na_v_1.1 may not be prominently expressed within MS ChAT neurons in our study: Na_v_1.1 and MS ChAT antibody staining were previously observed to colocalize, but that was in P60-180 rats (Bender et al., 2016), whereas our study focused upon P21-30 mice. Second, compensatory upregulation of other sodium channel subtypes (as in (Yu et al., 2006)) may have counteracted any functional reduction in sodium current in the *Scn1a*^+/-^ MS ChAT cells. Finally, MS ChAT cells, which have inherently slow-firing characteristics, may be less sensitive to changes in Na_v_1.1 expression levels, such that normal action potential generation can be maintained in these cells even with a reduction in sodium current.

Future studies could confirm Na_v_1.1 expression within the mouse MS ChAT neurons across different developmental timepoints, for instance using the recently generated *Scn1a*-GFP transgenic mouse line (Yamagata et al., 2023). It may also be the case that Na_v_1.1 has greater relevance for axonal propagation of action potentials than for action potential generation in MS ChAT cells (as in (Kaneko et al., 2022)); this could be explored by quantifying downstream cholinergic transmission within hippocampus. However, it is notable that hippocampal theta power was preserved in *Scn1a*^+/-^ mice (Jansen et al., 2021); as hippocampal theta power is mainly established by septal cholinergic inputs to hippocampus (Lee et al., 1994), this suggests that MS ChAT neurons are indeed likely to be functionally normal in this DS model.

### Diversity of firing patterns identified within MS PV neurons

Multiple prior studies have identified subtypes of MS neurons based upon their firing patterns (e.g. (Griffith, 1988; Jones et al., 1999; Mattis et al., 2014; Morris et al., 1999; Serafin et al., 1996; Sotty et al., 2003; Yi et al., 2021)). Our results further identify the diversity of firing patterns that can be seen even within the PV neuronal population.

Fast-firing neurons have been previously demonstrated to be PV-positive (Morris et al., 1999; Morris & Henderson, 2000; Yi et al., 2021). This is consistent with our data, especially noting that most WT MS PV neurons are in this category.

Burst-firing neurons, which fire only at the onset of a depolarizing pulse, have also been previously described in the MS (Jones et al., 1999; Morris et al., 1999; Sotty et al., 2003).

Although we did not test the voltage dependence of the burst response – which has been shown to be more pronounced when cells are hyperpolarized (Jones et al., 1999; Morris et al., 1999; Sotty et al., 2003) – our data demonstrate that at least a subset of burst-firing MS cells are PV-positive. A recent characterization of PV versus SST neurons in MS (Yi et al., 2021) did not identify this subtype – perhaps because it is less prominent in WT mice (**Figure 6**) – but this burst-firing pattern has been identified for PV neurons within other brain regions (e.g., (Brandenburg et al., 2021; Hughes et al., 2012; Qiu et al., 2024)).

Finally, cluster-firing cells have also been previously described in the MS (Mattis et al., 2014; Serafin et al., 1996; Sotty et al., 2003), and in one study several of these were found to be glutamatergic (Sotty et al., 2003). Our results suggest that MS PV cells also exhibit cluster firing properties, and indeed these cluster-firing cells had high maximal instantaneous firing frequency and narrow spike width (**Table 3**) characteristic of PV cells. However, we cannot exclude the alternative possibility that these cells were among those that were tdT-positive but PV-negative (**Figure 2C**). This could be clarified by biocytin-filling and post-staining these neurons, although the yield would be low as they constitute only ∼10% of those encountered with tdT-guided patching.

### MS PV neurons are impaired in *Scn1a*^*+/-*^ mice

We found that MS PV cells were in aggregate highly impaired in *Scn1a*^+/-^ mice (**Figure 2; Table 2**), and that this overall pattern was seen across subtypes of PV cells (**Figure 5; Table 3**). This is consistent with PV neuron impairment identified in other brain regions, including hippocampus and neocortex, in DS models (De Stasi et al., 2016; Favero et al., 2018; Mattis et al., 2022; Tai et al., 2014).

In addition to hypoexcitability within MS PV neuron subtypes in *Scn1a*^+/-^ mice, we also observed a genotype difference in the proportion of different PV subtypes present in the MS (**Figure 6**). The most straightforward explanation is differences in ion channel expression (e.g., (Chamberland et al., 2023; Qiu et al., 2024)) converting *Scn1a*^+/-^ PV cells from a fast-firing to burst-firing pattern. Additionally, neuromodulation can dynamically impact the firing pattern within individual inhibitory cells (e.g., (Goff et al., 2023; Goff & Goldberg, 2019; Prönneke et al., 2020)). Regardless of underlying mechanism, our results highlight that neuronal dysfunction in *Scn1a*^+/-^ mice may manifest not only as impaired firing within a given subtype of neurons, but also a shift in representation of different subtypes across the population.

What might be the circuit and behavioral implications of MS PV cell dysfunction in DS? Septal inhibitory neurons synapse selectively onto inhibitory neurons in hippocampus (Freund & Antal, 1988), thereby disinhibiting hippocampal principal cells (Tóth et al., 1997) and pacing hippocampal theta oscillations (Etter et al., 2023; Gerashchenko et al., 2001; Hangya et al., 2009). Reduced hippocampal theta frequency was observed in a haploinsufficient mouse model of DS (Jansen et al, 2021), which may contribute to the cognitive deficits seen in related mice (e.g. (S. Han et al., 2012; Ito et al., 2013)). WT rats with focal siRNA-induced knockdown of Na_v_1.1 exclusively within the MS also had reduced theta frequency (Bender et al., 2013, 2016), directly implicating the MS in this finding. Future studies will elucidate dysfunction of the septohippocampal circuit in *Scn1a*^+/-^ mice and will parse the relative contributions of septal and hippocampal circuit elements to impairment of theta oscillations and hippocampal-dependent cognitive tasks.

Therapeutic interventions to restore Na_v_1.1 expression levels have demonstrated tremendous preclinical promise in DS (Fadila et al., 2023; Z. Han et al., 2020; Mich et al., 2023; Mora-Jimenez et al., 2021; Tanenhaus et al., 2022; Yuan et al., 2023). Our results underscore the “whole-brain” nature of DS: optimizing these treatments to achieve a brain-wide rescue will likely maximize their therapeutic effect across both seizure and non-seizure symptoms of DS.

## Materials and methods

### Experimental animals

All animal use was conducted under protocols approved by the University of Michigan Institutional Animal Care and Use Committee (IACUC) and were in accordance with the National Institutes of Health (NIH) *Guide for the Care and Use of Laboratory Animals*. Mice were maintained in a controlled-temperature environment, on a 12-hour light/dark cycle, with access to food and water *ad libitum*. All mice were genotyped via PCR analysis of tissue obtained at P5-P10 via amputation of a toe at the most distal joint.

Mice carrying the *Scn1a*^*tm1Kea*^ targeted null allele on the 129S6/SvEvTac background (129S.*Scn1a*^*+/-*^, RRID:MMRRC_037107-JAX) were provided as a generous gift from the laboratory of Dr. Lori Isom. To generate mice used for this study, male 129S.*Scn1a*^*+/-*^ mice were crossed to either WT or Cre-driver female mice (see below) on the C57BL/6J (B6) background (RRID: IMSR_JAX:000664). Hemizygous progeny from these crosses (B6.*Scn1a*^*+/-*^) have a mixed 50:50 129S:B6 background and exhibit an overt phenotype including spontaneous seizures and premature lethality (Miller et al., 2014; Mistry et al., 2014).

To enable visualization of ChAT or PV neurons in *Scn1a*^*+/-*^ mice, we performed the following breeding to ultimately express tdT within these neuron types: Males homozygous for either the *Chat*^*tm2(cre)Lowl*^*/J* (RRID:IMSR_JAX:006410) or *Pvalb*^*tm1(cre)Arbr*^*/J* (RRID:IMSR_JAX:017320) transgenic Cre cassette were crossed with females homozygous for the Ai14 tdT reporter cassette (RRID:IMSR_JAX:007914), yielding double-heterozygous ChAT-Cre.tdT and PV-Cre.tdT mice. Progeny from this cross were bred with each other and their double-homozygous offspring were identified by PCR. Double-homozygous ChAT-Cre.tdT and

PV-Cre.tdT females were then crossed with 129S.*Scn1a*^*+/-*^ males and their progeny were used in all experiments.

### Acute slice preparation

Mice (P21-30) were deeply anesthetized using inhaled isoflurane, confirmed by a toe pinch test. Subsequently, their brains were swiftly removed and immersed in an ice-cold sucrose solution composed of the following concentrations (in mM): NaCl 87, sucrose 75, KCl 2.5, CaCl_2_ 1.0, MgSO_4_ 2.0, NaHCO_3_ 26, NaH_2_PO_4_ 1.25, and glucose 10, supplemented with 95% O_2_ and 5% CO_2_. Coronal brain slices of 300 μm were prepared utilizing a Leica VT-1200S vibratome (Leica Microsystems Inc., Buffalo Grove, IL, USA). These slices were transferred to a holding chamber filled with oxygenated artificial cerebrospinal fluid (ACSF) containing the following concentrations (in mM): NaCl 125, KCl 2.5, CaCl_2_ 2.0, MgSO_4_ 1.0, NaHCO_3_ 26, NaH_2_PO_4_ 1.25, and glucose 10. Following transfer, the slices were allowed to recover at 32°C for 30 minutes, followed by an additional 30 minutes at room temperature before initiating recording procedures.

### Slice electrophysiology

Whole-cell patch clamp recordings were conducted using a SliceScope Pro 6000 electrophysiology system (Scientifica). Slices were transferred to a recording chamber and continuously perfused with oxygenated artificial cerebrospinal fluid (ACSF), which was bubbled with 95% O_2_ and 5% CO_2_, maintaining a perfusion rate of approximately 1 mL/min. Slice physiology experiments were conducted at 31°C, with the exception of Hm1a wash-on experiments (see below). For whole-cell recordings, borosilicate glass electrodes were employed, pulled to achieve a tip resistance of 3-4 MΩ using a P-97 puller (Sutter Instruments). These electrodes were filled with a K-Gluconate internal solution (K-gluconate 130, KCl 6.3, EGTA 0.5, MgCl_2_ 1.0, HEPES 10, Mg-ATP 4.0, and Na-GTP 0.3, in mM). The pH of the internal solution was adjusted to 7.30 using KOH, and the osmolarity was set to 285 mOsm with 30% sucrose. Electrode positioning was facilitated using a PatchStar manipulator (Scientifica).

Current was injected as needed to maintain cells at -60 mV during the current step protocol. Throughout current clamp experiments, series resistance compensation (bridge balance) was consistently applied, with periodic readjustments as deemed necessary. No liquid junction potential correction was performed. Data were included for final analysis only from cells with a resting potential below -50 mV for ChAT cells and below -60 mV for PV cells.

Hm1a (Alomone Labs STH-601) was used at a 250 nM final concentration in ACSF. Hm1a was perfused at a rate of 1 mL/min after a baseline recording was obtained. To improve the stability of our recordings across longer time periods, we performed these experiments at room temperature and with a more limited range of current steps.

### Analysis of whole-cell electrophysiology data

The analysis was conducted blind to genotype using custom Python software (https://github.com/mattis-laboratory/Slice-Physiology-Automated-Analysis) and the pyABF package (available at https://pypi.org/project/pyabf). An action potential (AP) was defined as crossing 0 mV and having a threshold characterized by a derivative of the voltage (dV/dt) greater than 10 mV/ms. Single AP properties were derived from the initial AP elicited during the rheobase sweep, with rheobase defined as the minimal current injection necessary to evoke an AP using 1 s sweeps with 20 pA intervals. The current/frequency plot was generated based on the average steady-state firing frequency computed at each current step. Similarly, the fold- rheobase plot was derived from the same dataset, normalizing the current to each cell’s rheobase and considering only integer values of fold-rheobase. The maximal steady-state firing frequency was computed as the highest average firing frequency across all 1 s sweeps.

Additionally, the maximal instantaneous firing frequency was calculated as the inverse of the shortest inter-spike interval observed across all sweeps. Action potential duration (APD) 50 and 90 represented the durations from threshold to the points at which the AP achieved 50% or 90% of repolarization, respectively.

The inter-spike interval (ISI) covariation was calculated as the standard deviation divided by the mean of all ISIs in the maximum frequency sweep. To determine the length of the first burst in each maximum frequency sweep, we first determined the mean and standard deviation of all inter-spike intervals. We defined a burst as having terminated when the gap between APs was greater than the mean plus three standard deviations. The burst length was then the time difference from the last AP before the cessation and the first AP (Goff & Goldberg, 2019).

All other parameters were computed according to previously established methodologies (Mattis et al., 2022).

### Immunohistochemistry

Mice were deeply anesthetized with isoflurane and transcardially perfused with ice-cold PBS followed by 4% paraformaldehyde (PFA) in PBS. Brains were removed and post-fixed at 4°C overnight in 4% PFA, then equilibrated overnight in 30% sucrose at 4°C. Tissue was sectioned at a thickness of 40 μm using a frozen microtome (Leica Biosystems) and sections were placed in cryoprotectant (25% glycerol, 30% ethylene glycol in PBS, pH adjusted to 6.7 with HCl) for long-term storage at -20°C. Sections were washed in PBS prior to immunostaining to remove residual cryoprotectant.

For both ChAT and PV immunostaining, sections were blocked and permeabilized in PBS containing 0.3% Triton X-100 and 3% normal donkey serum or 10% normal goat serum as appropriate for the choice of secondary antibody. Sections were incubated for 24 or 48 hours at room temperature with guinea pig anti-PV (Synaptic Systems, cat. no. 195308) or rabbit anti-ChAT (Invitrogen, cat. no. PIPA529653), respectively. Primary antibodies were diluted 1:1000 in the appropriate blocking solution. Following primary antibody incubation, sections were washed with PBS. Fresh 1:500 dilutions of secondary antibodies in the appropriate blocking solution were prepared prior to incubation. Sections were incubated for 2 hours in goat anti-guinea pig Alexa Fluor 488 (ThermoFisher, cat no., A-11073) or for 3 hours in donkey anti-rabbit Alexa Fluor 488 (Jackson ImmunoResearch, cat. no. 711-545-152) for PV or ChAT immunostaining, respectively. All sections were then washed in PBS, incubated in 1:50,000 DAPI (ThermoFisher, cat. no. D1306) in PBS at room temperature for 15 minutes and washed a final time in PBS. Sections were mounted and cover-slipped with PVA-DABCO mounting medium (MilliporeSigma, cat. no. 10981). Images were acquired using a Nikon fluorescent microscope.

To label PNNs, free-floating sections were washed briefly in PBS. Sections were then incubated for 24 hours at room temperature with biotinylated *Wisteria floribunda* lectin (Vector Laboratories, cat. no. B-1355-2) diluted 1:1000 in PBS containing 0.3% Triton X-100 (PBST). Sections were then washed again with PBS and incubated for 2 hours at room temperature with streptavidin conjugated to Alexa Fluor 488 (ThermoFisher, cat. no. S11223) diluted 1:1000 in PBST. Sections were washed a final time with PBS, then mounted and cover-slipped with PVA-DABCO.

### Image Analysis

Intensity of fluorescence signal of labeled PNNs in MS was quantified using Nikon NIS-Elements software. Images of MS-containing sections stained for PNNs were acquired and a region of interest (ROI) was manually drawn around the MS. For each section, a second ROI was also drawn in an adjacent region containing no apparent fluorescent labeling to obtain a background signal for normalization. The Nikon software was used to measure the mean fluorescence intensity for each ROI; normalized mean fluorescence was then calculated by dividing the mean intensity of each MS-containing ROI by that of the background ROI within the same section. These data were analyzed and plotted using GraphPad Prism.

## Acknowledgments

We thank all members of the Mattis lab for helpful discussion. This work was supported by NIH NINDS K08 NS121464 (JM) and the Kenneth Eisenberg Taubman Emerging Scholar Award (JM).

## References

Bender, A. C., Luikart, B. W., & Lenck-Santini, P.-P. (2016). Cognitive Deficits Associated with Nav1.1 Alterations: Involvement of Neuronal Firing Dynamics and Oscillations. PLOS ONE, 11(3), e0151538. 10.1371/journal.pone.0151538

Bender, A. C., Natola, H., Holmes, G. L., Scott, R. C., & Lenck-Santini, P.-P. (2013). Focal Scn1a knockdown induces cognitive impairment without seizures. Neurobiology of Disease, 54, 297–307. 10.1016/j.nbd.2012.12.021

Brandenburg, C., Smith, L. A., Kilander, M. B. C., Bridi, M. S., Lin, Y.-C., Huang, S., & Blatt, G. J. (2021). Parvalbumin subtypes of cerebellar Purkinje cells contribute to differential intrinsic firing properties. Molecular and Cellular Neurosciences, 115, 103650. 10.1016/j.mcn.2021.103650

Buzsáki, G. (2002). Theta oscillations in the hippocampus. Neuron, 33(3), 325–340. 10.1016/s0896-6273(02)00586-x

Buzsáki, G., & Moser, E. I. (2013). Memory, navigation and theta rhythm in the hippocampal-entorhinal system. Nature Neuroscience, 16(2), 130–138. 10.1038/nn.3304

Chamberland, S., Nebet, E. R., Valero, M., Hanani, M., Egger, R., Larsen, S. B., Eyring, K. W., Buzsáki, G., & Tsien, R. W. (2023). Brief synaptic inhibition persistently interrupts firing of fast-spiking interneurons. Neuron, 111(8), 1264-1281.e5. 10.1016/j.neuron.2023.01.017

Chaunsali, L., Tewari, B. P., & Sontheimer, H. (2021). Perineuronal Net Dynamics in the Pathophysiology of Epilepsy. Epilepsy Currents, 21(4), 273–281. 10.1177/15357597211018688

Claes, L., Del-Favero, J., Ceulemans, B., Lagae, L., Van Broeckhoven, C., & De Jonghe, P. (2001). De Novo Mutations in the Sodium-Channel Gene SCN1A Cause Severe Myoclonic Epilepsy of Infancy. The American Journal of Human Genetics, 68(6), 1327– 1332. 10.1086/320609

Colgin, L. L. (2013). Mechanisms and Functions of Theta Rhythms. Annual Review of Neuroscience, 36(1), 295–312. 10.1146/annurev-neuro-062012-170330

Darra, F., Battaglia, D., Dravet, C., Patrini, M., Offredi, F., Chieffo, D., Piazza, E., Fontana, E., Olivieri, G., Turrini, I., Dalla Bernardina, B., Granata, T., & Ragona, F. (2019). Dravet syndrome: Early electroclinical findings and long-term outcome in adolescents and adults. Epilepsia, 60(S3), S49–S58. 10.1111/epi.16297

De Stasi, A. M., Farisello, P., Marcon, I., Cavallari, S., Forli, A., Vecchia, D., Losi, G., Mantegazza, M., Panzeri, S., Carmignoto, G., Bacci, A., & Fellin, T. (2016). Unaltered Network Activity and Interneuronal Firing During Spontaneous Cortical Dynamics In Vivo in a Mouse Model of Severe Myoclonic Epilepsy of Infancy. Cerebral Cortex, 26(4), 1778–1794. 10.1093/cercor/bhw002

Etter, G., van der Veldt, S., Choi, J., & Williams, S. (2023). Optogenetic frequency scrambling of hippocampal theta oscillations dissociates working memory retrieval from hippocampal spatiotemporal codes. Nature Communications, 14(1), 410. 10.1038/s41467-023-35825-5

Fadila, S., Beucher, B., Dopeso-Reyes, I. G., Mavashov, A., Brusel, M., Anderson, K., Ismeurt, C., Goldberg, E. M., Ricobaraza, A., Hernandez-Alcoceba, R., Kremer, E. J., & Rubinstein, M. (2023). Viral vector-mediated expression of NaV1.1, after seizure onset, reduces epilepsy in mice with Dravet syndrome. The Journal of Clinical Investigation, e159316. 10.1172/JCI159316

Favero, M., Sotuyo, N. P., Lopez, E., Kearney, J. A., & Goldberg, E. M. (2018). A Transient Developmental Window of Fast-Spiking Interneuron Dysfunction in a Mouse Model of Dravet Syndrome. The Journal of Neuroscience: The Official Journal of the Society for Neuroscience, 38(36), 7912–7927. 10.1523/JNEUROSCI.0193-18.2018

Freund, T. F., & Antal, M. (1988). GABA-containing neurons in the septum control inhibitory interneurons in the hippocampus. Nature, 336(6195), 170–173. 10.1038/336170a0

Gerashchenko, D., Salin-Pascual, R., & Shiromani, P. J. (2001). Effects of hypocretin–saporin injections into the medial septum on sleep and hippocampal theta. Brain Research, 913(1), 106–115. 10.1016/S0006-8993(01)02792-5

Goff, K. M., & Goldberg, E. M. (2019). Vasoactive intestinal peptide-expressing interneurons are impaired in a mouse model of Dravet syndrome. eLife, 8, e46846. 10.7554/eLife.46846

Goff, K. M., Liebergall, S. R., Jiang, E., Somarowthu, A., & Goldberg, E. M. (2023). VIP interneuron impairment promotes in vivo circuit dysfunction and autism-related behaviors in Dravet syndrome. Cell Reports, 42(6), 112628. 10.1016/j.celrep.2023.112628

Griffith, W. H. (1988). Membrane properties of cell types within guinea pig basal forebrain nuclei in vitro. Journal of Neurophysiology, 59(5), 1590–1612. 10.1152/jn.1988.59.5.1590

Griffith, W. H., & Matthews, R. T. (1986). Electrophysiology of AChE-positive neurons in basal forebrain slices. Neuroscience Letters, 71(2), 169–174. 10.1016/0304-3940(86)90553-7

Han, S., Tai, C., Westenbroek, R. E., Yu, F. H., Cheah, C. S., Potter, G. B., Rubenstein, J. L., Scheuer, T., de la Iglesia, H.O., & Catterall, W. A. (2012). Autistic-like behaviour in Scn1a+/− mice and rescue by enhanced GABA-mediated neurotransmission. Nature, 489(7416), 385–390. 10.1038/nature11356

Han, Z., Chen, C., Christiansen, A., Ji, S., Lin, Q., Anumonwo, C., Liu, C., Leiser, S. C., Meena, Aznarez, I., Liau, G., & Isom, L. L. (2020). Antisense oligonucleotides increase Scn1a expression and reduce seizures and SUDEP incidence in a mouse model of Dravet syndrome. Science Translational Medicine, 12(558). 10.1126/scitranslmed.aaz6100

Hangya, B., Borhegyi, Z., Szilágyi, N., Freund, T. F., & Varga, V. (2009). GABAergic Neurons of the Medial Septum Lead the Hippocampal Network during Theta Activity. The Journal of Neuroscience, 29(25), 8094–8102. 10.1523/JNEUROSCI.5665-08.2009

Hughes, D. I., Sikander, S., Kinnon, C. M., Boyle, K. A., Watanabe, M., Callister, R. J., & Graham, B. A. (2012). Morphological, neurochemical and electrophysiological features of parvalbumin-expressing cells: A likely source of axo-axonic inputs in the mouse spinal dorsal horn. The Journal of Physiology, 590(16), 3927–3951. 10.1113/jphysiol.2012.235655

Huh, C. Y. L., Goutagny, R., & Williams, S. (2010). Glutamatergic Neurons of the Mouse Medial Septum and Diagonal Band of Broca Synaptically Drive Hippocampal Pyramidal Cells: Relevance for Hippocampal Theta Rhythm. Journal of Neuroscience, 30(47), 15951– 15961. 10.1523/JNEUROSCI.3663-10.2010

Ito, S., Ogiwara, I., Yamada, K., Miyamoto, H., Hensch, T. K., Osawa, M., & Yamakawa, K. (2013). Mouse with Nav1.1 haploinsufficiency, a model for Dravet syndrome, exhibits lowered sociability and learning impairment. Neurobiology of Disease, 49, 29–40. 10.1016/j.nbd.2012.08.003

Jansen, N. A., Perez, C., Schenke, M., Beurden, A. W. van, Dehghani, A., Voskuyl, R. A., Thijs, R. D., Ullah, G., Maagdenberg, A.M.J.M. van den, & Tolner, E. A. (2021). Impaired θ-γ Coupling Indicates Inhibitory Dysfunction and Seizure Risk in a Dravet Syndrome Mouse Model. Journal of Neuroscience, 41(3), 524–537. 10.1523/JNEUROSCI.2132-20.2020

Jones, G. A., Norris, S. K., & Henderson, Z. (1999). Conduction velocities and membrane properties of different classes of rat septohippocampal neurons recorded in vitro. The Journal of Physiology, 517(Pt 3), 867–877. 10.1111/j.1469-7793.1999.0867s.x

Kaneko, K., Currin, C. B., Goff, K. M., Wengert, E. R., Somarowthu, A., Vogels, T. P., & Goldberg, E. M. (2022). Developmentally regulated impairment of parvalbumin interneuron synaptic transmission in an experimental model of Dravet syndrome. Cell Reports, 38(13), 110580. 10.1016/j.celrep.2022.110580

Lee, M. G., Chrobak, J. J., Sik, A., Wiley, R. G., & Buzsáki, G. (1994). Hippocampal theta activity following selective lesion of the septal cholinergic systeM. Neuroscience, 62(4), 1033–1047. 10.1016/0306-4522(94)90341-7

Licheni, S. H., Mcmahon, J. M., Schneider, A. L., Davey, M. J., & Scheffer, I. E. (2018). Sleep problems in Dravet syndrome: A modifiable comorbidity. Developmental Medicine & Child Neurology, 60(2), 192–198. 10.1111/dmcn.13601

Markram, H., & Segal, M. (1990). Electrophysiological characteristics of cholinergic and non-cholinergic neurons in the rat medial septum-diagonal band complex. Brain Research, 513(1), 171–174. 10.1016/0006-8993(90)91106-Q

Mattis, J., Brill, J., Evans, S., Lerner, T. N., Davidson, T. J., Hyun, M., Ramakrishnan, C., Deisseroth, K., & Huguenard, J. R. (2014). Frequency-Dependent, Cell Type-Divergent Signaling in the Hippocamposeptal Projection. Journal of Neuroscience, 34(35), 11769– 11780. 10.1523/JNEUROSCI.5188-13.2014

Mattis, J., Somarowthu, A., Goff, K. M., Jiang, E., Yom, J., Sotuyo, N., Mcgarry, L. M., Feng, H., Kaneko, K., & Goldberg, E. M. (2022). Corticohippocampal circuit dysfunction in a mouse model of Dravet syndrome. eLife, 11, e69293. 10.7554/eLife.69293

Mich, J. K., Ryu, J., Wei, A. D., Gore, B. B., Guo, R., Bard, A. M., Martinez, R. A., Bishaw, Y., Luber, E., Oliveira Santos, L. M., Miranda, N., Ramirez, J.-M., Ting, J. T., Lein, E. S., Levi, B. P., & Kalume, F. K. (2023). AAV-mediated interneuron-specific gene replacement for Dravet syndrome. bioRxiv, 2023.12.15.571820. 10.1101/2023.12.15.571820

Miller, A. R., Hawkins, N. A., McCollom, C. E., & Kearney, J. A. (2014). Mapping genetic modifiers of survival in a mouse model of Dravet syndrome. Genes, Brain, and Behavior, 13(2), 163–172. 10.1111/gbb.12099

Mistry, A. M., Thompson, C. H., Miller, A. R., Vanoye, C. G., George, A. L., & Kearney, J. A. (2014). Strain- and age-dependent hippocampal neuron sodium currents correlate with epilepsy severity in Dravet syndrome mice. Neurobiology of Disease, 65, 1–11. 10.1016/j.nbd.2014.01.006

Mora-Jimenez, L., Valencia, M., Sanchez-Carpintero, R., Tønnesen, J., Fadila, S., Rubinstein, M., Gonzalez-Aparicio, M., Bunuales, M., Fernandez-Pierola, E., Nicolas, M. J., Puerta, E., Miguelez, C., Minguez, P. G., Lumbreras, S., Gonzalez-Aseguinolaza, G., Ricobaraza, A., & Hernandez-Alcoceba, R. (2021). Transfer of SCN1A to the brain of adolescent mouse model of Dravet syndrome improves epileptic, motor, and behavioral manifestations. Molecular Therapy. Nucleic Acids, 25, 585–602. 10.1016/j.omtn.2021.08.003

Morris, N. P., Harris, S. J., & Henderson, Z. (1999). Parvalbumin-immunoreactive, fast-spiking neurons in the medial septum/diagonal band complex of the rat: Intracellular recordings in vitro. Neuroscience, 92(2), 589–600. 10.1016/S0306-4522(99)00026-3

Morris, N. P., & Henderson, Z. (2000). Perineuronal nets ensheath fast spiking, parvalbumin-immunoreactive neurons in the medial septum/diagonal band complex. European Journal of Neuroscience, 12(3), 828–838. 10.1046/j.1460-9568.2000.00970.x

Osteen, J. D., Herzig, V., Gilchrist, J., Emrick, J. J., Zhang, C., Wang, X., Castro, J., Garcia-Caraballo, S., Grundy, L., Rychkov, G. Y., Weyer, A. D., Dekan, Z., Undheim, E. A. B., Alewood, P., Stucky, C. L., Brierley, S. M., Basbaum, A. I., Bosmans, F., King, G. F., & Julius, D. (2016). Selective spider toxins reveal a role for Nav1.1 channel in mechanical pain. Nature, 534(7608), 494–499. 10.1038/nature17976

Osteen, J. D., Sampson, K., Iyer, V., Julius, D., & Bosmans, F. (2017). Pharmacology of the Nav1.1 domain IV voltage sensor reveals coupling between inactivation gating processes. Proceedings of the National Academy of Sciences of the United States of America, 114(26), 6836–6841. 10.1073/pnas.1621263114

Prönneke, A., Witte, M., Möck, M., & Staiger, J. F. (2020). Neuromodulation Leads to a Burst-Tonic Switch in a Subset of VIP Neurons in Mouse Primary Somatosensory (Barrel) Cortex. Cerebral Cortex, 30(2), 488–504. 10.1093/cercor/bhz102

Qiu, H., Miraucourt, L. S., Petitjean, H., Xu, M., Theriault, C., Davidova, A., Soubeyre, V., Poulen, G., Lonjon, N., Vachiery-Lahaye, F., Bauchet, L., Levesque-Damphousse, P., Estall, J. L., Bourinet, E., & Sharif-Naeini, R. (2024). Parvalbumin gates chronic pain through the modulation of firing patterns in inhibitory neurons. Proceedings of the National Academy of Sciences of the United States of America, 121(27), e2403777121. 10.1073/pnas.2403777121

Rubinstein, M., Han, S., Tai, C., Westenbroek, R. E., Hunker, A., Scheuer, T., & Catterall, W. A. (2015). Dissecting the phenotypes of Dravet syndrome by gene deletion. Brain, 138(8), 2219–2233. 10.1093/brain/awv142

Selvarajah, A., Gorodetsky, C., Marques, P., Zulfiqar Ali, Q., Berg, A. T., Fasano, A., & Andrade, D. M. (2022). Progressive Worsening of Gait and Motor Abnormalities in Older Adults With Dravet Syndrome. Neurology, 98(22), e2204–e2210. 10.1212/WNL.0000000000200341

Serafin, M., Williams, S., Khateb, A., Fort, P., & Mühlethaler, M. (1996). Rhythmic firing of medial septum non-cholinergic neurons. Neuroscience, 75(3), 671–675. 10.1016/0306-4522(96)00349-1

Sotty, F., Danik, M., Manseau, F., Laplante, F., Quirion, R., & Williams, S. (2003). Distinct electrophysiological properties of glutamatergic, cholinergic and GABAergic rat septohippocampal neurons: Novel implications for hippocampal rhythmicity. The Journal of Physiology, 551(Pt 3), 927–943. 10.1113/jphysiol.2003.046847

Tai, C., Abe, Y., Westenbroek, R. E., Scheuer, T., & Catterall, W. A. (2014). Impaired excitability of somatostatin- and parvalbumin-expressing cortical interneurons in a mouse model of Dravet syndrome. Proceedings of the National Academy of Sciences, 111(30), E3139– E3148. 10.1073/pnas.1411131111

Tanenhaus, A., Stowe, T., Young, A., McLaughlin, J., Aeran, R., Lin, I. W., Li, J., Hosur, R., Chen, M., Leedy, J., Chou, T., Pillay, S., Vila, M. C., Kearney, J. A., Moorhead, M., Belle, A., & Tagliatela, S. (2022). Cell-Selective Adeno-Associated Virus-Mediated SCN1A Gene Regulation Therapy Rescues Mortality and Seizure Phenotypes in a Dravet Syndrome Mouse Model and Is Well Tolerated in Nonhuman Primates. Human Gene Therapy, 33(11–12), 579–597. 10.1089/hum.2022.037

Tóth, K., Freund, T. F., & Miles, R. (1997). Disinhibition of rat hippocampal pyramidal cells by GABAergic afferents from the septum. The Journal of Physiology, 500(2), 463–474. 10.1113/jphysiol.1997.sp022033

Tsai, M.-S., Lee, M.-L., Chang, C.-Y., Fan, H.-H., Yu, I.-S., Chen, Y.-T., You, J.-Y., Chen, C.-Y., Chang, F.-C., Hsiao, J. H., Khorkova, O., Liou, H.-H., Yanagawa, Y., Lee, L.-J., & Lin, S.-W. (2015). Functional and structural deficits of the dentate gyrus network coincide with emerging spontaneous seizures in an Scn1a mutant Dravet Syndrome model during development. Neurobiology of Disease, 77, 35–48. 10.1016/j.nbd.2015.02.010

Vandecasteele, M., Varga, V., Berényi, A., Papp, E., Barthó, P., Venance, L., Freund, T. F., & Buzsáki, G. (2014). Optogenetic activation of septal cholinergic neurons suppresses sharp wave ripples and enhances theta oscillations in the hippocampus. Proceedings of the National Academy of Sciences, 111(37), 13535–13540. 10.1073/pnas.1411233111

Villas, N., Meskis, M. A., & Goodliffe, S. (2017). Dravet syndrome: Characteristics, comorbidities, and caregiver concerns. Epilepsy & Behavior, 74, 81–86. 10.1016/j.yebeh.2017.06.031

Wolff, M., Cassé-Perrot, C., & Dravet, C. (2006). Severe myoclonic epilepsy of infants (Dravet syndrome): Natural history and neuropsychological findings. Epilepsia, 47 Suppl 2. 10.1111/j.1528-1167.2006.00688.x

Yamagata, T., Ogiwara, I., Tatsukawa, T., Suzuki, T., Otsuka, Y., Imaeda, N., Mazaki, E., Inoue, I., Tokonami, N., Hibi, Y., Itohara, S., & Yamakawa, K. (2023). Scn1a-GFP transgenic mouse revealed Nav1.1 expression in neocortical pyramidal tract projection neurons. eLife, 12, e87495. 10.7554/eLife.87495

Yi, F., Garrett, T., Deisseroth, K., Haario, H., Stone, E., & Lawrence, J. J. (2021). Septohippocampal transmission from parvalbumin-positive neurons features rapid recovery from synaptic depression. Scientific Reports, 11(1), Article 1. 10.1038/s41598-020-80245-w

Yu, F. H., Mantegazza, M., Westenbroek, R. E., Robbins, C. A., Kalume, F., Burton, K. A., Spain, W. J., McKnight, G. S., Scheuer, T., & Catterall, W. A. (2006). Reduced sodium current in GABAergic interneurons in a mouse model of severe myoclonic epilepsy in infancy. Nature Neuroscience, 9(9), 1142–1149. 10.1038/nn1754

Yuan, Y., Lopez-Santiago, L., Denomme, N., Chen, C., O’Malley, H. A., Hodges, S. L., Ji, S., Han, Z., Christiansen, A., & Isom, L. L. (2023). ASO restores excitability, GABA signalling and sodium current density in a model of Dravet syndrome. Brain, awad349. 10.1093/brain/awad349

