## Supplemental figures and figure legends for "Medial septum parvalbumin-expressing inhibitory neurons are impaired in a mouse model of Dravet Syndrome"

**Supplementary figures and figure legends:**

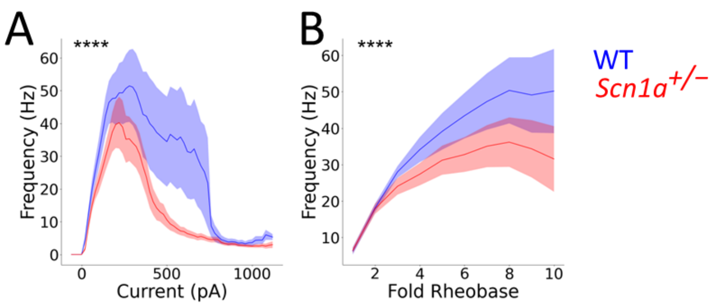


**Figure S1. MS ChAT-negative cells have impaired firing in *Scn1a*$^{+/-}$ mice.** tdT-negative (i.e., ChAT-negative) cells were patched within medial septum; these are expected to contain a mix of glutamatergic and GABAergic (PV and SST-expressing) neurons. Current/frequency (I-f) plot shows reduced firing in *Scn1a*$^{+/-}$ cells (red) compared to WT (blue). n = 16 cells / 7 mice for WT and n = 18 / 5 mice for *Scn1a*$^{+/-}$. Line and shaded areas represent mean and SEM. (****) indicates p < 0.0001; 2-way ANOVA.

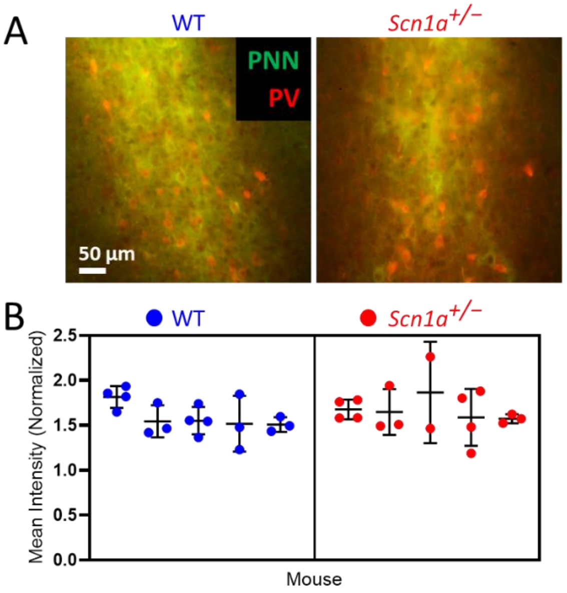


**Figure S2. Perineuronal nets in MS are comparable between WT and *Scn1a*$^{+/-}$ mice. (A)** Representative images of MS from WT (left) and *Scn1a*$^{+/-}$ (right) with immunostaining for PNNs (green) and PV (red). **(B)** Quantification of mean fluorescence intensity of PNN signal. 2-5 MS slices were quantified per animal. No significant difference was detected between genotypes (p = 0.49; nested t-test). n = 17 sections / 5 mice (WT) and 16 sections / 5 mice (*Scn1a*$^{+/}$). Cumulative intensity was also quantified and showed no significant difference between genotypes (data not shown).
